# Facial Emotions Perceptions in Young Female Students with High- Level Anxity and Subclinical Panic Disorder. ERP and BEhavioral Study

**DOI:** 10.1101/479899

**Authors:** V.Yu. Karpova, E.S. Mikhailova, N.Yu. Gerasimenko, A.B. Kushnir, S.A. Gordeev, N.N. Alipov

## Abstract

This study investigated facial emotion processing in non-medicated young students (girls) with panic disorder. 13 young girls with panic disorder and 14 matched healthy controls were recruited. Evoked potential (EP) components P100, N150, and P300 in the posterior areas, and N200, P300, and late negativity were evaluated while the participants recognize angry, fearful, happy, and neutral facial stimuli. The girls with panic disorder showed increased levels of situational anxiety compared to healthy controls. EP demonstrated an increased reactivity to facial expression at sensory stage (P100 component), in particular, on angry faces, that indicates a shift automated attention on threat facial stimuli. The increased reactivity was also found in later processing, corresponding to the P300 component, reflecting an enhanced selective attention to socially important events. In subjects with panic disorder, we also found signs of increased activation in the right temporal area in the P300 time window, and the increased late frontal negativity in 350-450 ms time window. It can be assumed, that altered functional state of the prefrontal regions results in reduced top-down modulating effects on the lower limbic and sensory cortex levels.

## INTRODUCTION

The ability to identify and interpret facial expressions of emotion is crucial to functioning in social networks and to interpersonal relationships. This is because facial expressions can signal our emotional states and those of others, and also because they influence the production and regulation of affective states and behavior according to these signals (Phillips et al., 2003). P. Ekman (Ekman, 1992) has described six, which are often called the six ‘basic’ emotions. These include sadness, happiness, surprise, anger, disgust, and fear. Affective states may then influence sensitivity to and selective attention towards facial emotional expressions. Ekman’s studies report that many populations (including isolated tribes) are able to recognize facial expressions of these emotions. Abnormal facial emotion processing has been studied in a range of psychiatric conditions. This is of interest, firstly because studies of facial emotion processing may provide important information regarding abnormalities of regional brain functioning in psychiatric patients. Secondly, abnormal facial emotion processing may directly give rise to some of the affective and social symptoms in subjects with psychiatric disorders. Thirdly, changes in facial emotion processing may help in the prediction of or monitoring of response to treatment in major depression (Bourke et al., 2010).

Today’s world with its stream of negative information and stressing factors creates strong prerequisites for neurotic disorders development. According to current data, the prevalence of anxiety disorders varies from 13.6 to 28.8% (Kessler et al., 2009). Such a high frequency of anxiety, notably in young, urges the search of neurophysiologic correlates of anxious states. It was shown that the anxiety, as well as the depression, influences the recognition of facial expression (Langenecker et al., 2005; Garner et al., 2009). In studies with eye movements tracking, for instance, patients with a generalized anxiety disorder, unlike healthy subjects, fixed the gaze primarily on stimuli associated with a threat (Mogg et al., 2000).

In neurological practice, non-epileptic paroxysmal disorders are widespread and panic disorder is one of them. Their main manifestation are repeated paroxysms of anxiety - panic attacks, which are not limited to a certain situation or circumstances and are therefore unpredictable. Patients with panic disorder are extremely sensitive to dangerous environmental signals and are highly sensitive to unexpected anxiety events, which is one of the key moments in maintaining the disease. Common symptoms are sudden heartbeats, chest pains, choking, dizziness and a sense of unreality. In a panic attack, patients often experience dramatically increasing fear and vegetative symptoms. The subjects also reported the following symptoms, manifested during periods of emotional stress (for example, preparation for exams): sweating, numbness of limbs, blanching, convulsions in the extremities, tremor, insomnia and fear of death.

Studies on people with anxiety disorder and panic attacks established impairment in emotion recognition and problems with communication (Windmann et al., 2002; Lange et al., 2012; Cao et al., 2017; Reutter et al., 2017). According to the literature, patients with anxiety disorders are characterized by a shift in attention to external stimuli that signal a potential threat, which is reflected in the parameters of cortical potentials (Windmann et al., 2002; Reutter et al., 2017).

In modern psychiatry and neurology, there are clear criteria for the separation of patients and healthy people. Most of the biomedical research is directed to a comparative analysis of such outlined groups. Much less attention is paid to the analysis of that part of the spectrum of disorders where the manifesting forms of diseases are presented. The search for biological markers of the initial stages of diseases promotes early detection, timely intervention, prevention of serious neurological and psychiatric disorders and is an important component of preventive medicine.

In the present work we investigated the behavioral characteristics and cortical visual event-related potentials (ERPs) in the task of recognizing individuals with different emotional expressions: reporting danger (anger and fear), positive (joy) and emotionally neutral in young girls with non-medicated subclinical panic disorder with rare panic attacks (all are students of the Pirogov Russian National Research Medical University, Moscow, Russia).

## METHODS

### Subjects

Thirteen untreated patients (females, 19.8±0.6 years) with panic attack (PA) and 14 (females, 21.5±0.5 years) healthy subjects (H) were recruited for this study from the Pirogov Russian National Research Medical University. In the PA patients diagnoses were based on the Structured Clinical Interview for Diagnostic and Statistical Manual of Mental Disorders, 4th edition (DSM-IV). Patients were excluded if they had diseases of the central nervous system, medical histories of alcohol or drug abuse, experience with electrical therapy, mental retardation, or a history of head injuries with loss of consciousness. They received no medication and did not attend any rehabilitation or psychotherapy courses. All subjects were right-handed. The study was performed accordingly to the ethical principles for medical research involving human subjects of the World Medical Association Declaration of Helsinki. Each patient and healthy subject was informed of the aims of the study and provided signed informed consent. All subjects were tested using Spilberg’s questionnaire on anxiety state (State-Trait Anxiety Inventory, STAI). Investigations were carried out in the first half of the day from 9 a.m. till 2 p.m.

### Stimuli

Stimuli were color photos of emotional faces taken from standardized Radboud University database (Langner et al., 2010). The angular size of the stimulus was 13 degrees vertically and 8 horizontally. Ultimately, there were 256 stimuli (8 for each emotion, 4 male and 4 female – fig. 1).

**Fig. 1.**
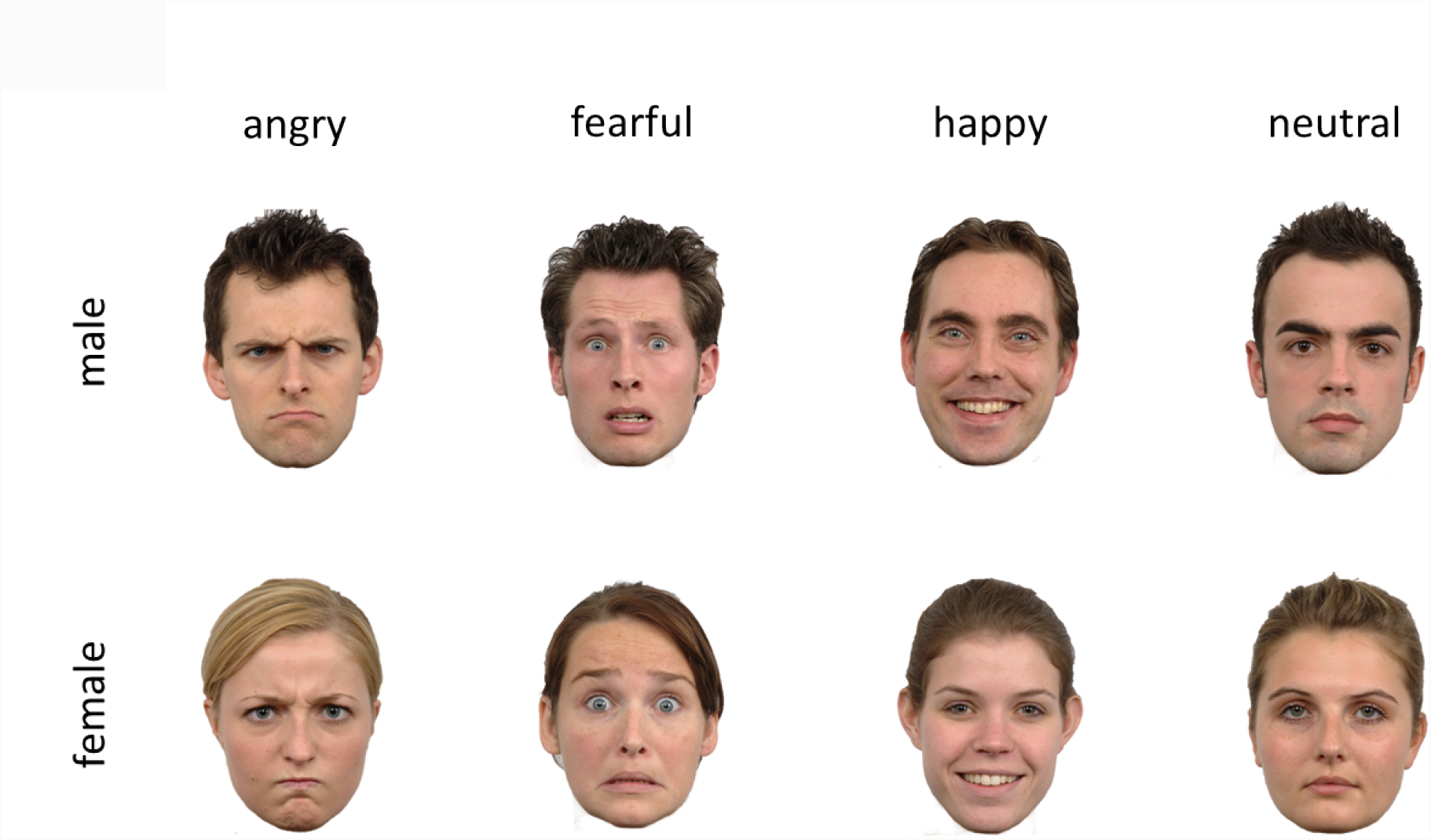
Stimuli: examples of faces.

**Fig. 2.**
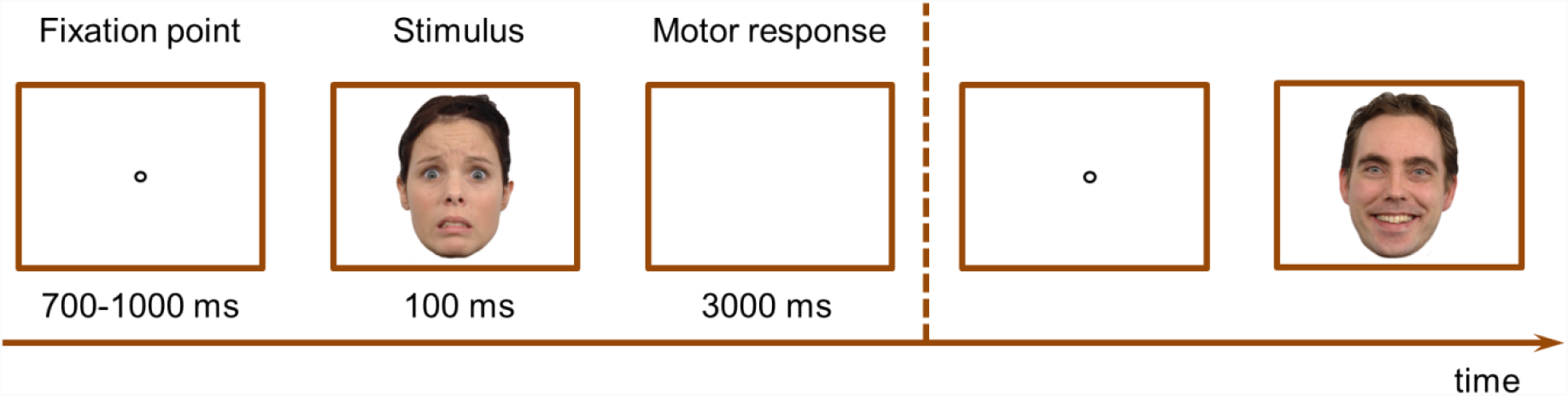
Procedure.

### Procedure

During the experiment the participants sat in the comfortable armchair in dark and sound-isolated room at a distance 120 centimeters from the computer monitor (19” NEC MultiSync EA193Mi, 1280 × 1024 pixel resolution, 60 Hz refresh rate). They were instructed to keep their head and eyes oriented towards fixation throughout testing and to blink only between trials. Stimuli were presented in the center of the monitor screen by E-Prime 2.0 program (Psychology Software Tools, Inc., USA). Every trial included fixation point (lengths varying between 700– 1000 ms), facial stimulus presentation (100ms) and delay (3000 ms). Every type of emotional facial expression (angry, fear, happy and neutral) was displayed on the computer screen 64 times, and the total number of stimuli was 256. The order of different emotions presentation was pseudo-random.

Participants were instructed to recognize facial expression shown on the screen and press corresponding bottom on the E-Prime 2.0 keyboard (Psychology Software Tools, Inc.). Before the session the stimuli were shown, and participants were instructed which button corresponds to which particular expression. Subjects were instructed to maintain fixation for the entire experiment and to press a button as fast as possible at the termination of each stimulation period, to maintain attention. The individual accuracy (probability of correct answers in %) and RT (ms) were averaged separately for each orientation across all session. The trials involving incorrect or missing responses were excluded from the analysis of RT. Before the session all participants completed a practice block of 20 trials.

### EEG recording and signal processing

Electroencephalography was recorded on EEG recording system with 128 channels Geodesics system (*HydroCel Geodesic Sensor Net,* Electrical Geodesics Inc., USA) that matched the head size of the participant. Electrode impedances were adjusted to stabilize under 50 kΩ before recording, and continuous EEG was sampled at 500 Hz with filters 0.01 to 100 Hz using the vertex (electrode № 129) as reference, and then was re-referenced off-line to the averaged reference and filtered using a band pass of 0.3 – 45 Hz.

The off-line analysis of the EEG data was undertaken using the Net Station 4.5.4 EEG Software. Notch filters were employed to remove electrical noise from the mains power supply (50 Hz) and the stimulus display monitor (60 Hz refresh rate). The data were segmented relative to the stimulus onset (300 ms before and 1000 ms after) and were averaged for each emotion category. The trials with blinks, saccades and non-ocular artifacts (epochs beyond the criteria of ±80 µV in any channel) were rejected from further analysis. The artifact-free EEGs were then averaged for each subject across all correct trials separately for each emotional expression. The minimum number of epochs included in any individual average was greater than 30. A final result was obtained by averaging across the subject averages for each emotional expression. The baseline, that began at 300 ms pre-stimulus and lasted for 300 ms, was used to perform a baseline correction

### ERP analysis

The peak amplitude of distinct ERP components were included into our analysis: in the occipital and temporal areas P100 (60-140 ms time window), N150 (115-200 ms), P300 (200-265 ms) peaks and for frontal areas N100 (70-130 ms), P150 (120-180 ms), N200 (170-260 ms) and P300 (220-320 ms) peaks. In the individual ERPs the average amplitude value (adaptive maximum/minimum) were measured within the time segments which corresponded to the analyzed components. The adaptive maximum/minimum for the components was counted within the 4-ms interval (2 ms before and 2 ms after the component peak). Additionally, based on the visual inspection, amplitudes in anterior late negativity component (350-600ms time window) were also analyzed. For each subject, a matrix was obtained with averaged ERPs for each emotion. Then, in the MatLab programming environment, a function was set that calculated the average amplitude at a given segment in intervals of 50 ms for certain electrodes (in the left frontal group for №№ 21, 22, 25 and the right frontal group for №№ 14, 9, 8).

The data obtained from individual electrodes at posterior and frontal regions were averaged to create groups of electrodes that covered occipital, temporal and frontal scalp regions over the left and right hemispheres (Fig. 3). The individual electrodes that were included in these groups were: the left occipital group - 66, 70 (O1), 71); the right occipital group - 76, 83 (O2), 84; the left temporal group - 50, 57, 64; the right temporal group - 100, 101, 95; the left frontal group - 21, 22 (FP1), 25 and the right frontal group -14, 9 (FP2), 8.

**Fig. 3.**
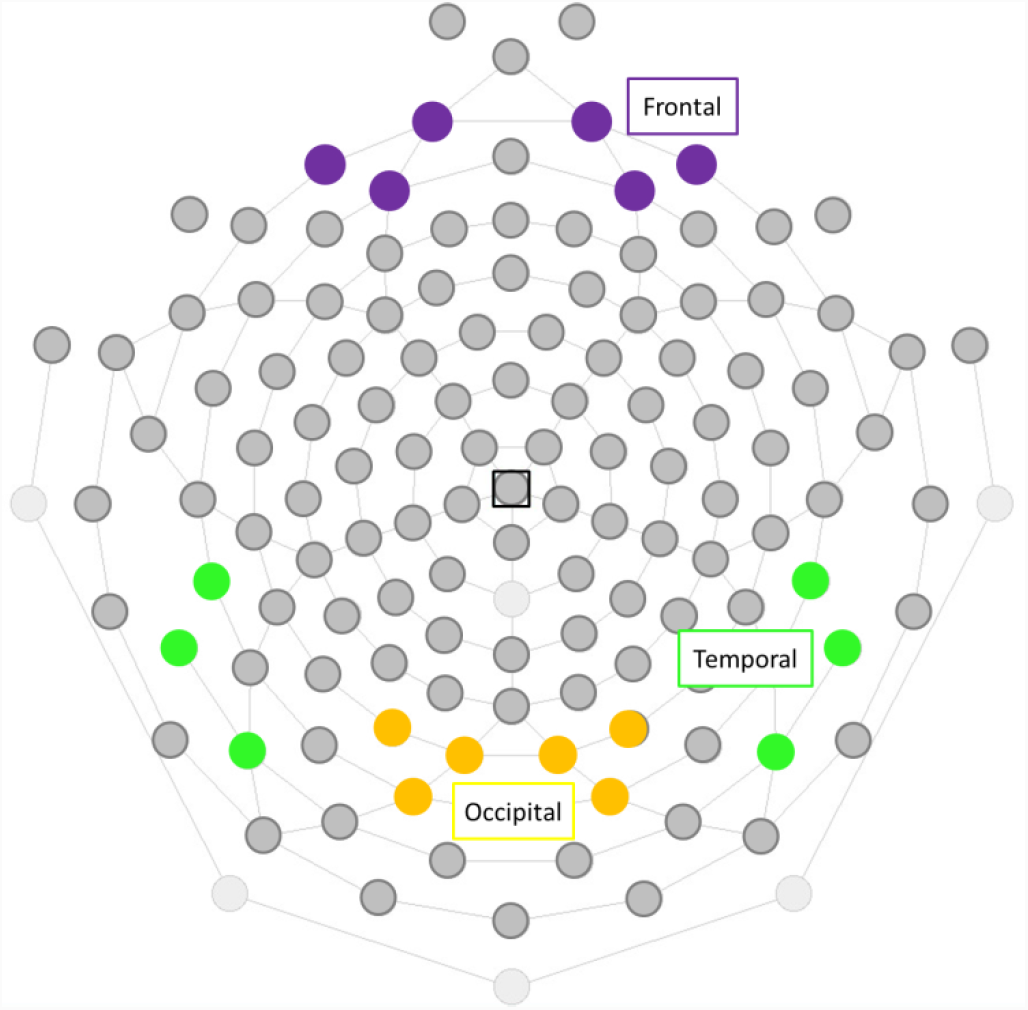
Arrangement of the high-density electrode arrays and the locations of the groups of electrodes over the skull (128 electrodes, EGI Geodesics) + key brain areas. Electrode grouping for the symmetrical occipital groups (yellow), for the temporal groups (green), and for the frontal groups (purple).

### Statistical analysis

The amplitude values of the ERP components were analyzed using a repeated measures analysis of variance (ANOVA RM) with Emotion (angry, fear, happiness, neutral expression), Hemisphere (the right and left) as within-subjects variable and Group as between-subjects variable. Psychometric data was analyzed using ANOVA RM with Emotion as within-subjects variable and Group as between-subjects variable. The Greenhouse-Geisser correction for nonsphericity was applied whenever appropriate. Significant main effects and interactions were further analyzed with post-hoc Newman Keuls Multiple Range Tests (alpha 0.05).

### Modeling of spreading dipole sources of activity

The modeling was carried out in the time window of the P100 (60–140 ms) and P300 (220-320 ms) components using the MatLab-based package Brainstorm 3.2. (Biomedical Imaging Group). We used the wMNE (L2-weightened Minimum Norm Estimates) method with the ICBM 152 standard anatomy, the OpenMEEG BEM realistic three-dimensional head model, and the standard electrode coordinates corresponding to the ASA-10-20 scheme. Three-dimensional spreading sources (spreading of activation through the brain volume) were reconstructed together with their projection on the cortical surface (surface spreading sources). The modeling was performed using group-averaged EPs for each of the four facial expressions for correct responses. A current source density distribution was obtained for the time intervals corresponding to the development of the P100 and P300 components of EPs in the occipital and temporal regions. To correlate the modeling results with anatomical brain structures, surface spreading sources were projected to the regions of interest (ROIs) of the Desikan–Killiany atlas.

## RESULTS

### Anxiety scores

The girls with PA differed from healthy subjects by higher values of personal anxiety (42.92 ± vs 57.27 ± 4.47, p = 0.02, T-test). Intergroup differences on situational anxiety were not found (36.00 ± 2.38 vs 36.91 ± 4.56).

### Behavioral data

For reaction time ANOVA RM revealed a significant main effect for the Emotion [F3.72=10.46; p=0.00001], Group [F_1,24_=7.13; p=0.01] and interaction effect of Emotion × Group [F_3,72_=3.08; p=0.03]. Healthy girls did not demonstrate any significant differences between emotions. In contrast, the girls with PA showed more high RT for negative emotional expressions compared with happy and neutral ones (0.0001<p<0.02, Neuman-Keuls). Mean RT in healthy and PA subjects were: for anger 780.54 ± 39.92 vs 907.69 ± 54.15 ms, accordingly; for fear 780.11 ± 34.22 and 965.40 ± 73.65 ms; for happy 713.92 ± 30.62 and 776.00 ± 31.67 ms; for neutral expression 741.64 ± 34.89 and 856.95 ± 49.00 ms. Generally, the girls with PA demonstrated higher RT compared to healthy subjects, which corresponds to a significant Group effect, but we did not find any significant between-group differences when every emotion was analyzed separately.

For accuracy ANOVA RM revealed only significant main effect for Emotion [F_3,75_=3.52; p=0.02]. We found no significant between-group differences.

### Analysis of ERPs

Statistical analyses of the ERP data were targeted at examining emotional expression effects on the ERPs at occipital, temporal and frontal electrode sites. Initially, repeated measure ANOVA RM (Area× Hemisphere ×Emotion ×Group) on the P100, N150 and P300 amplitudes performed over caudal groups of electrodes, specifically occipital and temporal groups. The analysis calculated for P100 amplitude revealed significant effects of Area [F_1,25_=80.16; p<0.0001], and significant interactions Area × Group [F_1,25_=7.56; p=0.01], Area × Emotion × Group [F_3,75_=3.06; p=0.03] and Area × Hemisphere × Group [F_3,75_=6.14; p=0.02].

Similarly, ANOVA RM on N150 and P300 amplitudes revealed the effect of Area and significant interactions of Area and other factors. For N150 main effect of Area was found [F_1,25_=13.23; p=0.001]. For P300 ANOVA RM revealed main effect of Area [F_1,23_=68.68; p<0.0001] and interactions: Area × Emotion × Group [F_3,69_=9.49; p<0.0001] and Area × Emotion × Hemisphere ×Group [F_3,69_=3.83; p<0.03]. Consequently, Area strongly influences the effects of other factors. For more precise assessment of between-group differences in the occipital and temporal areas, we conducted ANOVA RM separately in occipital and temporal groups of electrodes.

### Occipital group of electrodes P100 component

ANOVA RM revealed significant main effects of Group [F_1,25_=8.017; p=0.01]. The PAsubjects demonstrate the higher P100 amplitude compared to healthy ones (Fig. 7).

**Fig. 6.**
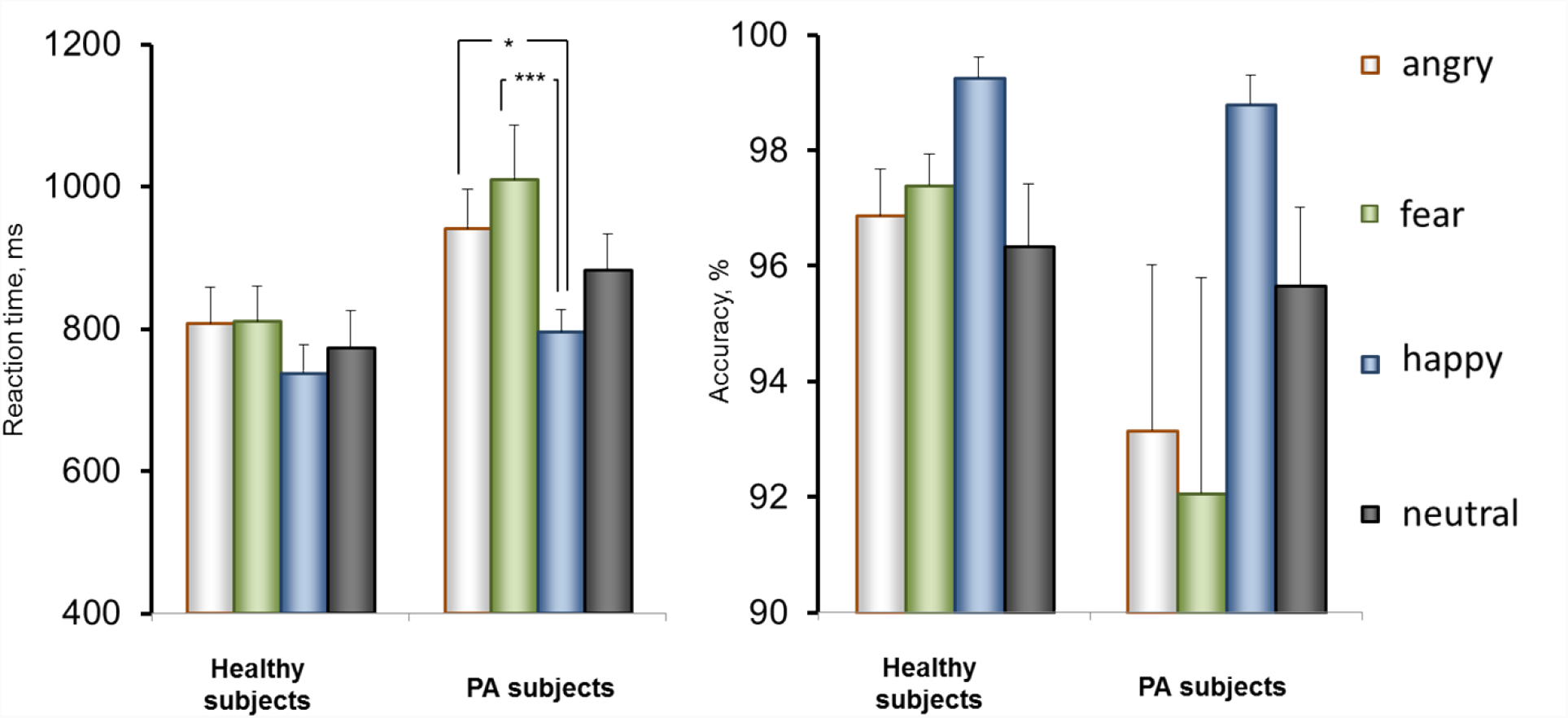
Bar charts for reaction time and accuracy for Healthy and PA subjects. *Error bars* denote standard error. \**p* < 0.05.

**Fig. 7.**
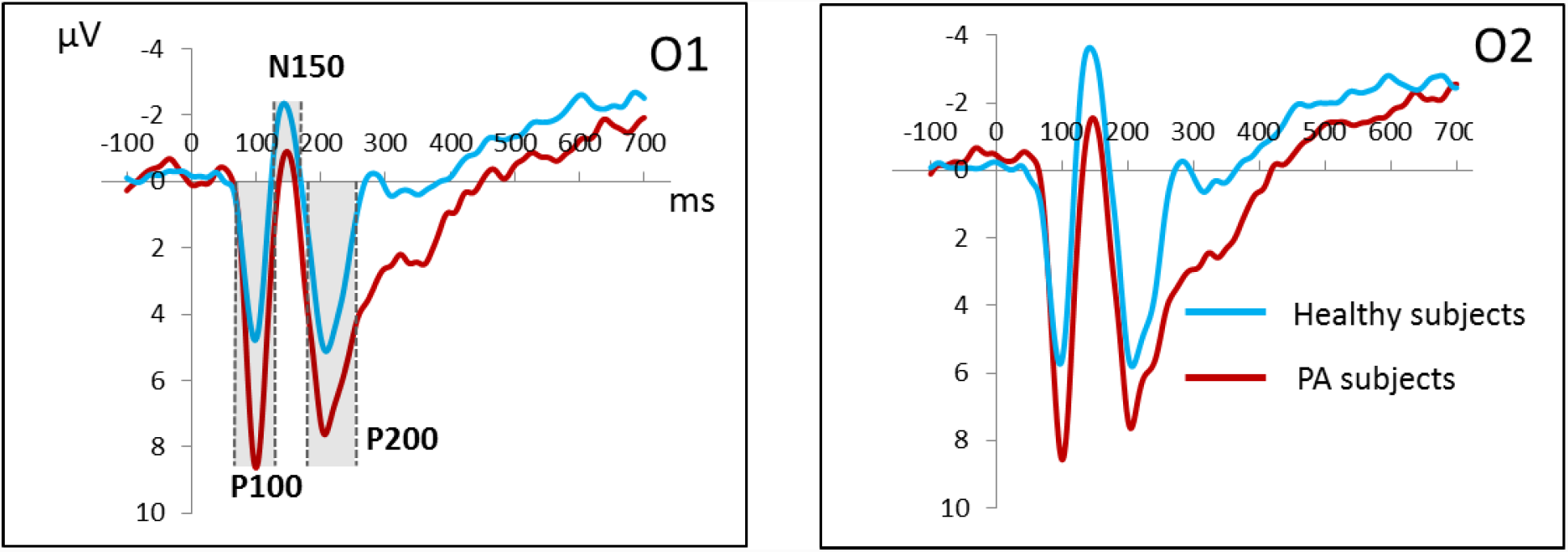
Grand average ERP waveforms for angry facial expressions at the occipital (O1 and O2) electrodes in the healthy group (blue) and PA group (red). Vertical scale represents voltage amplitude in μV and horizontal scale displays latency in ms.

Significant between-group differences were found over all emotions. In the left hemisphere the differences were found to be more pronounced than the ones in the right one (0.005<p<0.01 and 0.01<p<0.05, accordingly). Furthermore, in the left hemisphere negative emotions (angry, fear) showed more significant between-groups differences compared with the differences elicited by happy and neutral ones (p<0.005 *vs* p<0.01, accordingly).

Taking into account the significant Group effect, further we examined emotion recognition in each of groups. Healthy subjects were not found to have any significant differences in recognition of various emotions. In contrast, in PA subjects ANOVA RM (Emotion × Hemisphere) showed a significant effect of Emotion [F_3,36_=3.6; p = 0.02]. Follow up comparisons revealed higher ?100 amplitude for angry than for other facial expressions (p<0.05, Neuman-Keuls) in PA group (Fig. 7).

### N150 component

ANOVA RM on amplitude of N150 revealed significant main effects of Emotion [F_3,75_=2.62; p=0.05], Hemisphere [F_1,25_=11.19; p=0.003], and interactions Emotion x Hemisphere [F_3,75_=4.36; p=0.01]. Further paired comparisons showed that the amplitude of N150 is higher in the right hemisphere than in the left for fearful (p=0.0002) and happy (p=0.006) expressions, that corresponds to significant effect of Hemisphere and Emotion × Hemisphere interactions. For Group no significant interaction or main effects were found.

### P300 component

There was significant main effect of Group [F_1,24_=5.87; p=0.02], that manifested in the greater P300 amplitude in PA subjects compared with healthy control (Fig. 9). Also, main effects of Emotion [F_3,72_=3.86; p=0.01] and interaction Emotion x Hemisphere [F_3,72_=4.33; p=0.007] were found.

**Fig. 8.**
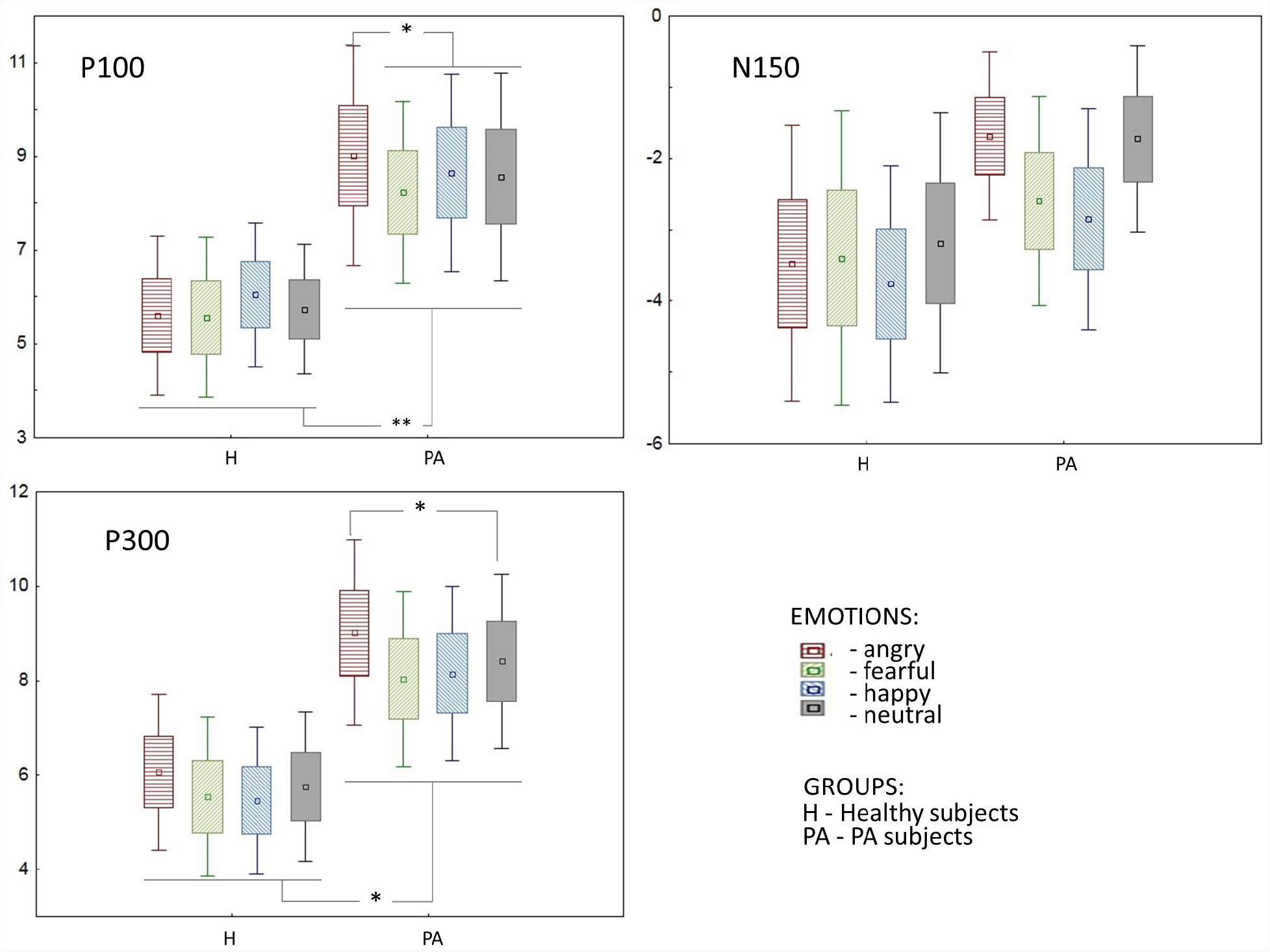
Box plots for components P100, N150 and P300 in the right occipital electrode group. Mean ±SE. Wiskers represent 0.95 conf. interval.

**Fig. 9.**
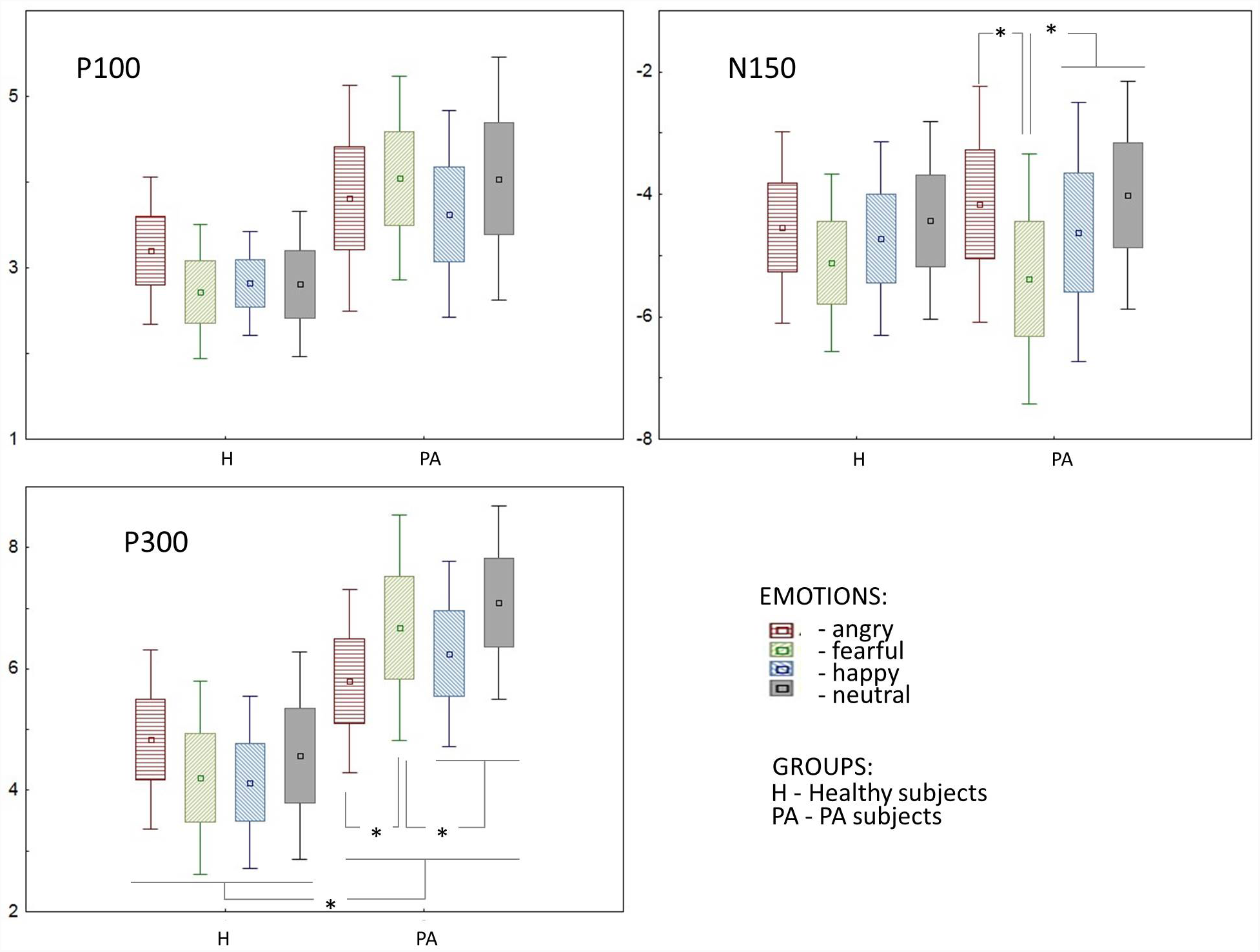
Box plots for components P100, N150 and P300 in the right temporal electrode group. Mean ±SE. Wiskers represent 0.95 conf. interval.

Taking into account the significant effect of Group, ANOVA RM analysis was performed in PA and healthy subjects separately. PA subjects showed significant effect of Emotion [F_3,36_=3.54; p=0.02] and interaction Emotion × Hemisphere [F_3,36_=4.84; p=0.006]. Further post-hoc comparisons showed larger P300 amplitudes for angry faces compared with fearful (p=0.02), happy (p=0.02) and neutral (p=0.05) ones in the right hemisphere, and for angry faces compared with neutral one (p=0.01) in the left hemisphere. No significant effects and interactions were found in the healthy controls.

### Temporal group of electrodes

#### P100 component

There was significant effect of Hemisphere [F_1,25_=8.55; p=0.007], while the effects of Group and Emotion were insignificant. P100 amplitude in the right temporal group of electrodes were greater than in the left one (total for all emotions 4.49±0.44 *vs* 3.50±0.31; T=2.83, df=26, p=0.01).

#### N150 component

ANOVA RM on the N150 amplitude did not reveal a significant main effect of Group suggesting the proximity of N150 amplitudes in the healthy and PA subjects (Fig. 9 and Fig. 10). However, there were significant effects of Emotion [F_3,75_=11.28; p=0.0001], Hemisphere [F_1,25_=9.92; p=0.05], and interaction Emotion × Hemisphere [F_3,75_=4.68; p=0.005] with the greater N150 on fearful faces compared with other ones in the right hemisphere (0.0001<p<0.003), (Fig. 9).

**Fig. 10.**
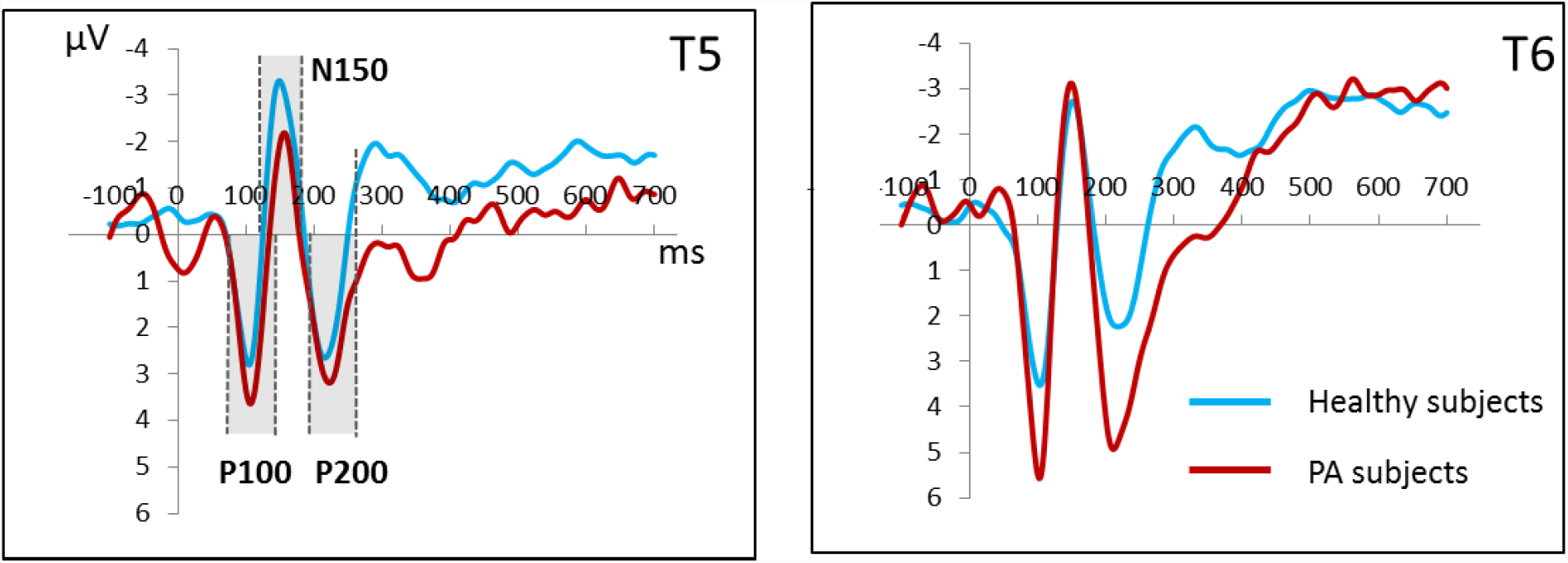
Grand average ERP waveforms for angry facial expressions at the temporal (T5 and T6) electrodes in the healthy group (blue) and group with panic attacks (red). Vertical scale represents voltage amplitude in μV and horizontal scale displays latency in ms.

#### P300 component

The amplitude of the later component P300 is higher in the PA group, as evidenced by the significant Group effect (F_1,24_ = 6.43; p=0.05). Also, a significant interaction Emotion × Hemisphere × Group (F_3,72_ = 3.89; p=0.05) was found. Further, ANOVA RM conducted separately in groups of healthy and PA subjects. Significant effects were found only in the PA group. There was found an interaction Emotion × Hemisphere (F_3,36_ = 5.78; p=0.01), which appeared in higher amplitude of P300 component for frightened and neutral faces compared to angry and happy (0.01<p<0.05) only in the right hemisphere. A comparison of the P300 component in the two hemispheres showed a greater P300 amplitude in the right hemisphere compared to the left, but it was significant (p <0.05) only for emotionally neutral faces.

#### Frontal group of electrodes

When analyzing the amplitude of ERP components, ANOVA RM (Emotion × Hemisphere × Group) revealed significant effects only for N200 and P300. Below we present these results.

#### N200 component

ANOVA RM on the N200 amplitude reveal a significant main effect of Group [F_1,25_=4.35; p=0.05]. As shown in Fig. 11 and 12, the main effect of Group manifested in greater N200 amplitude in PA subjects compared to healthy controls. Also, there were a significant interaction Emotion × Hemisphere [F_3,75_=3.3; p=0.025], and an effect of Emotion that did not reach significance level [F_3,75_=2.80; p=0.09].

**Fig. 11.**
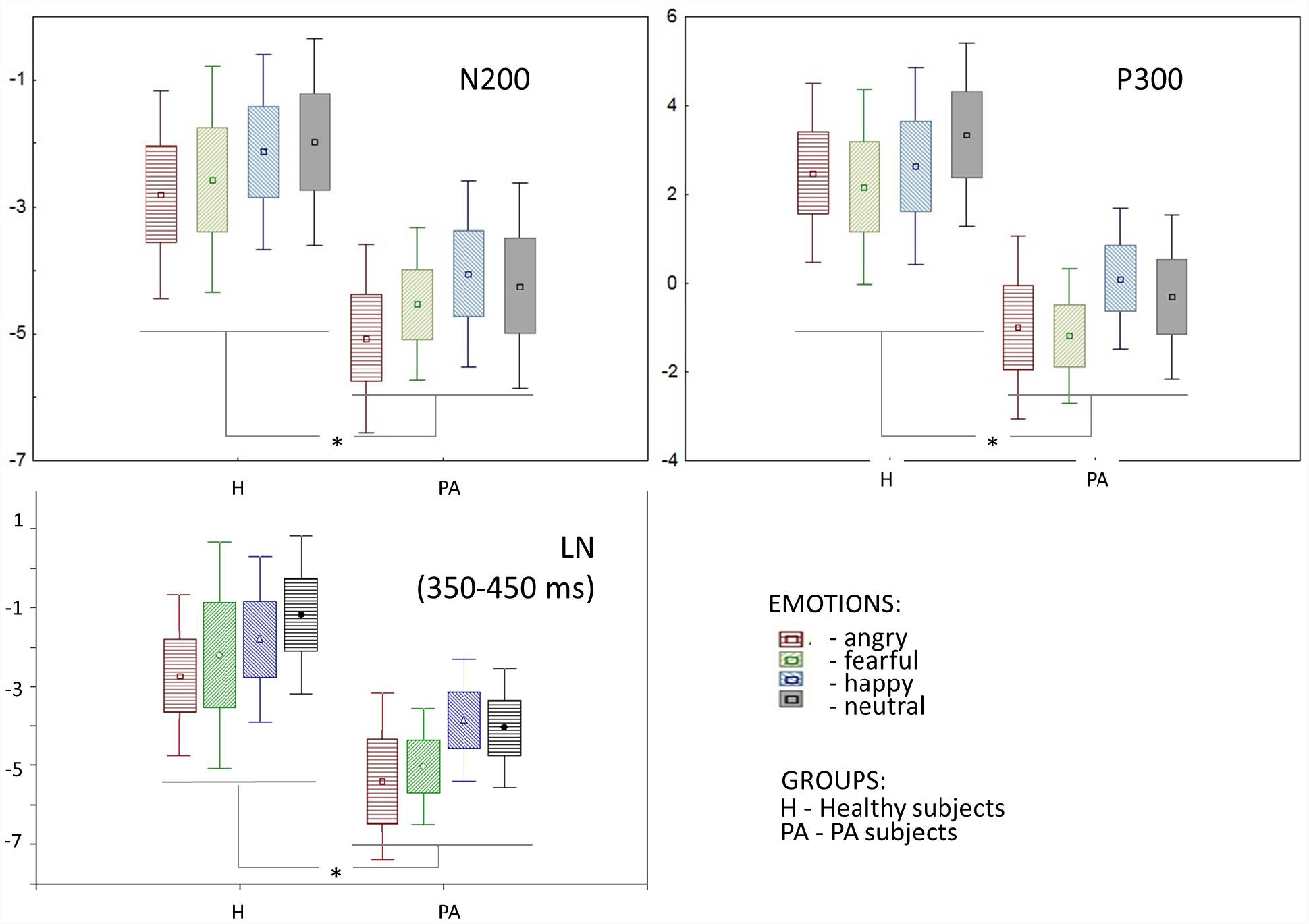
Box plots for amplitude of N200 and P300 and LN mean amplitude in 350-450 ms time period in the right frontal electrode group. Mean ±SE. Wiskers represent 0.95 conf. interval.

**Fig. 12.**
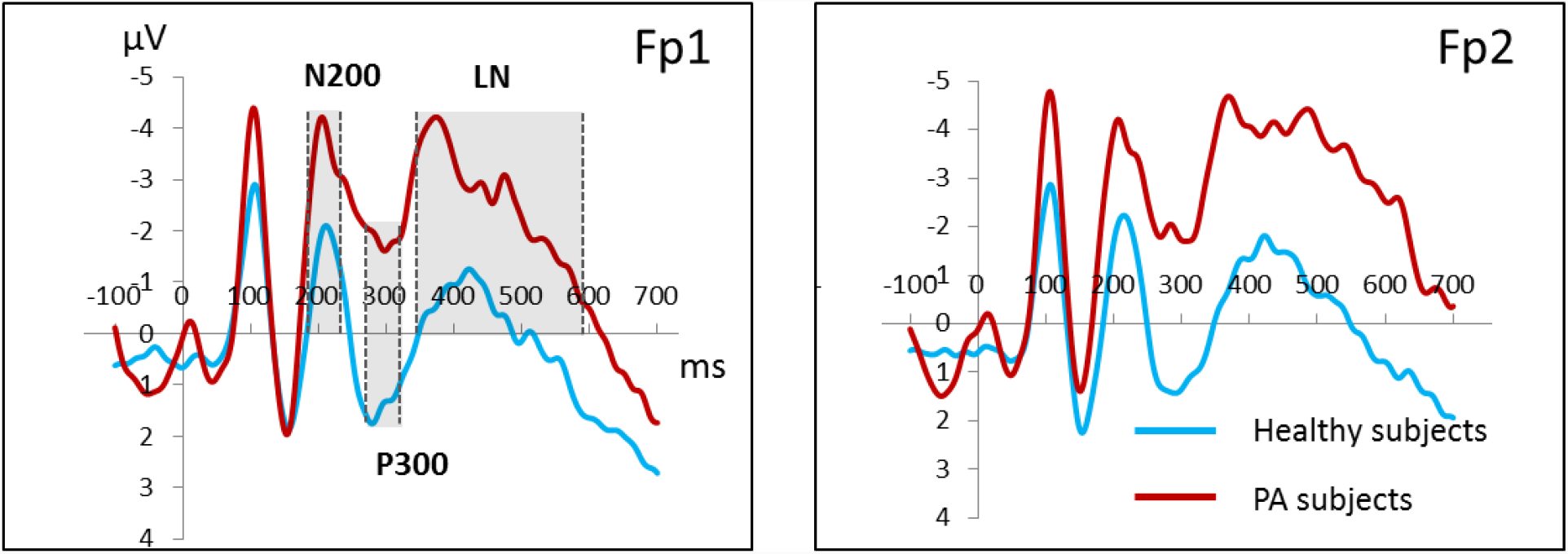
Grand average ERP waveforms for angry facial expressions at the frontal (Fp1 and Fp2) electrodes in the healthy group (blue) and PA group (red). Vertical scale represents voltage amplitude in μV and horizontal scale displays latency in ms.

**Fig. 13.**
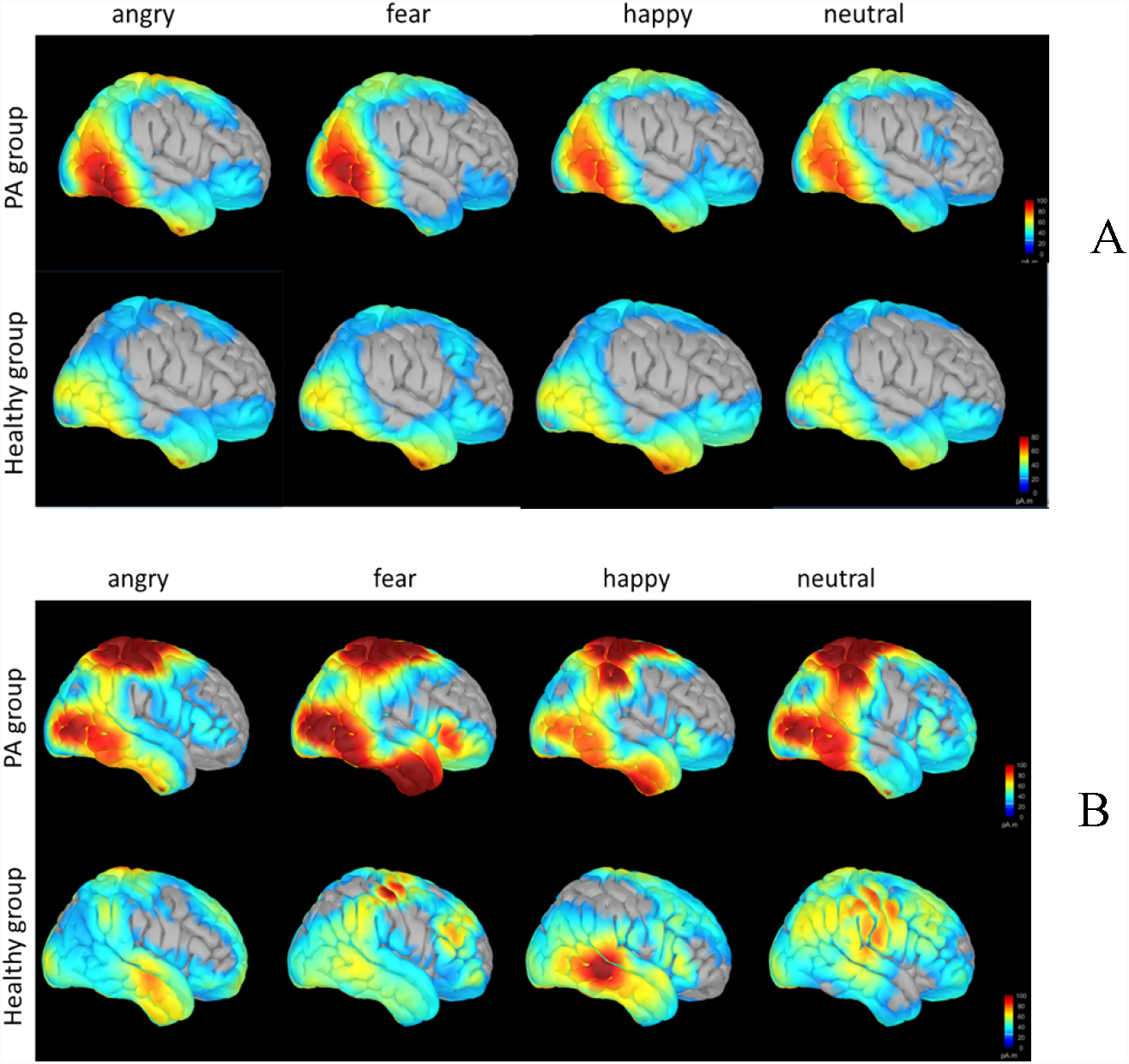
The modeling of the distributed sources of the P100 and P300 components by the wMNE method. Surface distributed sources for fearful, angry, happy and neutral expressions in healthy and PA groups. A - modeling of the P100, time - 104 ms after the stimulus. B - modeling of the P300, time 220 ms after the stimulus.

In the healthy subjects follow-up analysis revealed a significant interaction Emotion × Hemisphere [F_3,39_=3.77; p=0.02]. But the analysis of contrasts did not reveal any significant differences between the emotions. In PA subjects ANOVA RM revealed no significant effects.

#### P300 component

ANOVA RM on the P300 amplitude reveal a significant main effect of Group [F_1,25_=6.43; p=0.02]. Also, there were a significant effect of Emotion [F_3,75_=3.18; p=0.02] and an interaction Emotion × Hemisphere [F_3,75_=2.86; p=0.04]. As shown in Fig. 12, the P300 amplitude is more negative in PA group than in healthy subjects. A follow-up analysis did not reveal any significant effects and interactions in both groups.

#### Mean LN amplitude in interval 350-450 ms

ANOVA RM on mean amplitude reveal significant main effects of Group [F_1,23_=5.29; p=0.03] and Emotion [F_3,33_=2.73; p=0.06]. As shown in Fig. 12, the amplitude is greater in PA group than in healthy subjects. Taking into account main effect of Group, ANOVA RM analysis was performed in PA and healthy subjects separately. PA subjects demonstrated an effect of Emotion [F_3,33_=2.73; p=0.06]. Post-hoc analysis revealed the greater LN amplitude for angry faces compared with happy and neutral ones (p=0.05) only in the right hemisphere. The healthy subjects demonstrated an effect of Emotion [F_3,33_=2.73; p=0.06]. In the right hemisphere angry faces evoked the greater LN amplitude compared with neutral ones (p=0.02).

## DISCUSSION

We compared psychometric and neurophysiological correlates of facial emotional expression in girls with panic attacks and healthy subjects. The two groups were taken from the same social group (2nd year students of the medical academy), they were about the same age, and girls with panic attacks did not receive medical treatment. The girls of the PA group had infrequent seizures, mainly due to situations of emotional stress during the examination session. They differed from healthy people with an increased level of situational anxiety, and demonstrated the differences in psychometric indicators of the identification of emotions. In the PA group, negative facial expressions linked to anxiety (anger and fear) were identified with a large RT and a large number of errors compared to happy and emotionally neutral facial expressions. In the healthy control, such differences were absent. These data correspond to the literature about the perception impairment of negative emotional facial expressions in subjects with an increased level of anxiety (Bourke et al., 2010; Wieser et al., 2012; Langeslag, van Strien, 2017).

In the PA group, changes in the processing of information about facial expression were revealed by indicators of evoked activity, which were recorded at various stages of processing and at different cortical levels from the sensory zones to the prefrontal cortex. Following our analysis, we found the increased P100 amplitude in the occipital cortex in PA subjects, which may be an indicator of increased involuntary attention to the facial pattern, which signals about possible threat or its absence, and affects social behavior. At the same time, the higher reactivity of P100 component to angry faces among PA subjects indicates a higher level of involuntary attention to threatening faces. According to the literature, panic and anxiety disorders are associated with early affective pre-attentive evaluation of the stimulus (Windmann, 1998). That is associated with the automatic unconscious processing of this information, as reflected in early changes in the EP obtained during our study. These results are also consistent with the results of Rossignol and coauthors (Rossignol et al., 2012) who described the enhanced perceptual responses during visual processing of facial stimuli in young socially anxious individuals but without panic attack symptoms.

Similar between-group differences were found for later the P300 positivity, which is a cognitive decision-related component of the ERP, associated also with selective attention (Donchin et al., 1973). The higher P300 amplitude observed in the PA subjects indicates an increased level of selective attention to the facial pattern, its “grabbing” from the visual environment. As a consequence, the increased influence of information about faces on the social behavior of people with PA. Furthermore, the enlarged P300 amplitude suggested that the facial expression of anger is particularly meaningful for the PA subjects. This result indicates that the P300 amplitude reflects the processing of information-rich events and corresponds to the large number of studies that emphasize the relationship between the P300 and information content (Kok, 2001; Bledowski et al., 2004; Ferrari et al., 2010). Also, these results are consistent with the results of Wieser and coauthors (Wieser et al., 2012), where angry and neutral facial stimuli were used as distractors in the task of orientation recognizing. It has been shown that in the socially anxious individuals the threatening facial expressions are preferentially processed and involuntarily capture and divert attention resources from simultaneous demands.

In the temporal area, which is closely related to the processing of faces and facial expression, between-group differences were found only for the P300 component, which was higher in the PA group. At the same time, the increased amplitude of the earlier P100 and N150 components in the right hemisphere was found both in PA subjects and in healthy controls. The main feature of the ERP in the temporal cortex is an increase of the P300 amplitude in girls with PA and the presence of pronounced between-hemisphere asymmetry of P300 component in them with a predominance of activity on the right one. P300 differences between PA and healthy control are higher in the right hemisphere. In a study by Besteher and coauthors (Besteher et al., 2017), the authors found that the gray matter increase in the middle and a superior temporal and inferior parietal gyrus is observed only in individuals with subclinical anxious.

Also, the PA group demonstrated an increased contribution of negativity in the frontal areas, which manifested itself in the form of an increase of the amplitude of N200, a negative shift of the P300 component and an increase of the amplitude of late slow negativity in the range of 350-450 ms. In literature the effects in the time window between 300 and 500 ms post-stimulus are considered as an index of unconscious memory and automatic familiarity processes (Paller, 2000). The timing of the differences between the two groups (between 350 and 450 ms) suggests that they are related to automatic memory and familiarity processes more than to consciously controlled ones. The similar differences were found by Windmann and coauthors (Windmann et al., 2002), when they compared the patients with panic disorder and healthy controls performing an old/new recognition memory task with emotionally negative and neutral words.

Consequently, in the PA subjects, the executive structures of the frontal cortex, along with the sensory and limbic areas, are involved in a circuit of structures that respond with enhanced answer to faces, especially signaling of danger. It corresponds to the opinion, that in case of depression and anxiety, failure of prefrontal regions to flexibly modulate and inhibit emotional reactions and evaluations engendered by these limbic structures may cause the affective biases and cognitive abnormalities that have been described for patients with anxiety and depression (Windmann et al., 2002; Cao et al., 2017; Stephanou et al., 2018).

Overall, in our study, it was shown that girls from urban subclinical populations (students) with increased levels of situational anxiety and subclinical panic disorder differ from healthy controls by increased reactivity to the facial expression already at sensory processing stage in the early visual cortex, that indicates a shift automatic attention on faces, in particular, on ones signaling the threat. Increased facial cortical reactivity was also found at the later stage of facial processing, corresponding to the P300 component, reflecting, as well as the P100, enhanced attention to socially important events. In girls with panic disorder we also found signs of heightened activation of the temporal area, i.e. in limbic structures, which were more pronounced in the right hemisphere, and increased the late frontal negativity (350-450 ms poststmulus window). It can be assumed, that altered functional state of the prefrontal regions results in reduced top-down modulating effects on the limbic and sensory cortex levels. In general, the girls from urban population suffering from the subclinical form of panic disorder demonstrate the increased vigilance and reactivity of sensory and limbic structures to facial stimuli, especially signaling about the danger.

## Acknowledgment.

The study was supported by Russian Foundation for Basic Research grant No. 16-06-00945-OGN, «Factors of psychosocial disadaptation at persons with various forms psychovegetative disorders».

## REFERENCES

1. Besteher, B., Gaser, C., Langbein, K., Dietzek, M., Sauer, H., & Nenadić, I. (2017). Effects of subclinical depression, anxiety and somatization on brain structure in healthy subjects. J Affect Disord, 215, 111–117.

2. Bledowski, C., Prvulovic, C, D., Hoechstetter, K., Scherg, M., Wibral, M., Goebel, R., & Linden, D.E. (2004). Localizing P300 generators in visual target and distractor processing: a combined eventrelated potential and functional magnetic resonance imaging study. J Neurosci, 24, 9353–9360.

3. Bourke, C., Douglas, K., & R. Porter, R. (2010). Processing of facial emotion expression in major depression: a review. Aust N Z J Psychiatry, 44, 681–696.

4. Cao, J., Liu, Q., Li, Y., Yang, J., Gu, R., Liang, J., Qi, Y., Wu, H., & Liu, X. (2017) Cognitive behavioural therapy attenuates the enhanced early facial stimuli processing in social anxiety disorders: an ERP investigation. Behav Brain Funct, 13, 12.

5. Donchin, E., Kubovy, M., Kutas, M., Johnson, R., & Herning, R.I. (1973). Graded changes in evoked response (P300) amplitude as a function of cognitive activity. Percept Psychophys, 14, 319–324.

6. Ekman, P. (1992). Facial expressions of emotion: new findings, new questions. Psychol Sci, 3, 34–38.

7. Ferrari, V., Bradley, M.M., Codispoti, M., & Lang, P.J. (2010). Detecting novelty and significance. J Cogn Neurosci, 22, 404–411.

8. Garner, M., Baldwin, D., Bradley, B., & Mogg, K. (2009). Impaired identification of fearful faces in Generalised Social Phobia. J Affect Disord,115, 460–465.

9. Kessler, R. C., Aguilar-Gaxiola, S., Alonso, J., Chatterji, S., Lee, S., Ormel, J., Ustün, T. B., & Wang, P, S. (2009). The global burden of mental disorders: An update from the WHO World Mental Health (WMH) surveys. Epidemiol Psichiatry Soc, 18, 23–33.

10. Kok, A. (2001). On the utility of P3 amplitude as a measure of processing capacity. Psychophysiology, 38, 557–577.

11. Lange, W. G., Allart, E., Keijsers, G. P, Rinck, M., & Becker, E. S. (2012). A neutral face is not neutral even if you have not seen it: social anxiety disorder and affective priming with facial expressions. Cogn Behav Ther, 41, 108–118.

12. Langenecker, S. A., Bieliauskas, L. A., Rapport, L. J., Zubieta, J. K., Wilde, E. A., & Berent, S. (2005). Face emotion perception and executive functioning deficits in depression. J Clin Exp Neuropsychol, 27, 320–333.

13. Langeslag, S. J. E., & van Strien, J. W. (2018). Early visual processing of snakes and angry faces: An ERP study. Brain Res, 1678, 297–303.

14. Langner, O., Dotsch, R., Bijlstra, G., Wigboldus, D. H. J., Hawk, S. T., & van Knippenberg A. (2010). Presentation and validation of the Radboud Faces Database. Cogn Emot, 24, 1377—1388.

15. Mogg, K., Millar, N., & Bradley, B. P. (2000). Biases in eye movements to threatening facial expressions in generalized anxiety disorder and depressive disorder. J Abnorm Psychol, 109, 695–704.

16. Paller, K. A. (2000). Neural measures of conscious and unconscious memory. Behavioral Neurology, 12, 127–141.

17. Phillips, M. L., Drevets, W.C., Rauch, S. L., & Lane, R. (2003). Neurobiology of emotion perception I: the neural basis of normal emotion perception. Biol Psychiatry, 54, 504–514.

18. Reutter, M., Hewig, J., Wieser, M. J., & Osinsky, R. (2017). The N2pc component reliably captures attentional bias in social anxiety. Psychophysiology, 54, 519–527.

19. Rossignol, M., Campanella, S., Maurage, P., Heeren, A., Falbo, L., Philippot, P. (2012). Enhanced perceptual responses during visual processing of facial stimuli in young socially anxious individuals. Neurosci. Lett, 526, 68–73.

20. Stephanou, K., Davey, C. G., Kerestes, R., Whittle, S., & Harrison, B. J. Hard to look on the bright side: neural correlates of impaired emotion regulation in depressed youth. Soc Cogn Affect Neurosci, 12, 1138–1148.

21. Wieser, M. J., McTeague, L. M., & Keil, A. (2012). Competition effects of threatening faces in social anxiety. Emotion, 12, 1050–1060.

22. Windmann, S. (1998). Panic disorder from a monistic perspective: integrating neurobiological and psychological approaches. J Anxiety Disord, 12, 485–507.

23. Windmann, S., Sakhavat, Z., & Kutas, M. (2002). Electrophysiological evidence reveals affective evaluation deficits early in stimulus processing in patients with panic disorder. J Abnorm Psychol, 111, 357–369.

